# A negatively charged unstructured loop autoinhibits mammalian Dicer and supports fidelity of miRNA biogenesis

**DOI:** 10.64898/2026.02.19.706781

**Authors:** Diego F. Joseph, Nikola Noskova, Radek Malik, David Zapletal, Josef Pasulka, Valeria Buccheri, David Buchta, Jan Petrovsky, Karel Kubicek, Petr Svoboda, Richard Stefl

## Abstract

Dicer RNases produce small RNAs for RNA interference (RNAi) and microRNA (miRNA) pathways. Although vertebrate Dicers share a conserved overall architecture with other eukaryotic Dicers, they are uniquely adapted for the production of gene-regulating miRNAs. We report here this adaptation extends beyond the structured core of the enzyme and relies on an intrinsically disordered region (IDR) conserved across jawed vertebrates. This negatively charged IDR supports fidelity of miRNA biogenesis while inhibiting RNAi. The IDR projects into a positively charged substrate-binding groove of Dicer and stabilizes a pre-dicing state, which effectively licenses authentic miRNA precursors. Electron cryo-microscopy shows that removal of the IDR shifts the equilibrium from the pre-dicing to the dicing state, thus increasing enzymatic activity and broadening substrate accessibility. In cells, deletion, replacement or variations of the IDR affect fidelity of microRNA biogenesis, change substrate specificity, and enhance RNAi. Altogether, our work highlights functional relevance of unstructured protein parts, which escape attention in structural analyses, and introduces an autoinhibitory IDR element mimicking negative RNA charge as a component of the molecular grammar governing IDRs in RNA-binding proteins.

## INTRODUCTION

Dicer endoribonucleases make small RNAs in miRNA and RNAi pathways, fundamental sequence-specific silencing mechanisms in eukaryotes. (Zapletal *et al*, 2023). A typical Dicer architecture resembles the letter “L”, with the PAZ domain at the top, tandem RNase III domains in the central catalytic core, and a complex helicase domain forming the base (Lau *et al*, 2012; Lau *et al*, 2009; Liu *et al*, 2018; Taylor *et al*, 2013). The helicase domain has a clamp-like architecture of the RIG-I family of RNA helicases (Fairman-Williams *et al*, 2010) and is composed of three globular subdomains: an N-terminal DExD/H subdomain (HEL1), which is separated by an insertion subdomain (HEL2i) from a helicase superfamily C-terminal subdomain (HEL2).

The helicase domain is the key structural element segregating miRNA from the RNAi pathways *in vivo*. Multifunctional RNAi of invertebrates (Ketting, 2011) relies on processive cleavage of long double-stranded RNA (dsRNA), which enters Dicer via the helicase in an ATP-dependent manner (Cenik *et al*, 2011; Ketting *et al*, 2001; Liu *et al*, 2003; Welker *et al*, 2011) and is cleaved into small interfering RNAs (siRNA).

In contrast, biogenesis of gene-regulating miRNAs comprises a single, precise cleavage of genome-encoded small stem-loop precursors (pre-miRNA) (reviewed in (Bartel, 2018)), which does not require the helicase activity. During miRNA biogenesis, the PAZ domain binds the 3′ overhang of a pre-miRNA hairpin, while the helicase domain contacts the apical pre-miRNA loop, thereby bridging the two distal arms of the “L”-shaped Dicer, forming triangular RNA–protein assembly. This triangular RNA–protein assembly establishes a cleavage-incompetent checkpoint (“pre-dicing state”) that discriminates authentic pre-miRNAs from other RNA molecules. Subsequently, Dicer switches into a cleave-competent state, which entails repositioning of the substrate into the RNA processing center where the two RNase III domains function as catalytic “half sites”, each cleaving one strand of the double-stranded substrate (MacRae *et al*, 2006; Zhang *et al*, 2004).

Mammalian Dicers mainly produce miRNAs. Although their HEL1 domain has invariantly conserved residues that would be necessary for ATPase activity (Cordin *et al*, 2006; Jia *et al*, 2017), the helicase domain does not exhibit the ATPase activity (Provost *et al*, 2002; Zhang *et al*, 2002), inhibits RNAi (Kennedy *et al*, 2015; Ma *et al*, 2008), and is critical for fidelity of miRNA biogenesis. (Zapletal *et al*, 2022). The HEL1 domain locks the mammalian Dicer in the closed “L”-shaped conformation to facilitate pre-miRNA recognition. Subsequent transition to the cleavage-competent state (“dicing” state) involves disengagement of the helicase domain from the mammalian Dicer core and the unlocked helicase becomes highly flexible (invisible in cryo-EM) (Lee *et al*, 2023; Zapletal *et al*., 2022). This differs from miRNA biogenesis in *Drosophila* where Dicer-1 carries a degenerated helicase domain, which remains in the cleavage-competent state fully bound to the Dicer core and undergoes only a small conformational change, which is sufficient to accommodate authentic miRNA precursors (Jouravleva *et al*, 2022).

We turned attention to the function of intrinsically disordered regions (IDRs), which have been invisible in structural analyses and remained the least understood parts of Dicer. Inspection of experimentally determined mammalian Dicer structures (Lee *et al*., 2023; Zapletal *et al*., 2022) and their comparison with AlphaFold predictions (Abramson *et al*, 2024; Jumper *et al*, 2021) has revealed four major IDRs forming loops at conserved positions. We report here that the first of the four IDRs, the negatively charged IDR1, functions as an autoinhibitory element acting in concert with the HEL1 domain to suppress RNAi and supports high-fidelity miRNA biogenesis.

## RESULTS & DISCUSSION

### Negatively charged IDR 1 is predicted to bind into the substrate channel

IDR1 originates from the helicase domain and carries a strong negative charge. AlphaFold 3 predictions insert IDR1 into the positively charged substrate channel of Dicer (Fig. 1A and S1A). The sequence with the negative charge on IDR1 is highly conserved across jawed vertebrates (Fig. 1B and S1B), which emerged over 400 million years ago (Zhu *et al*, 2013). Sequence, size, charge and predicted positions of IDR1 of earlier branching chordates and deuterostomes vary (Fig. 1B). For instance, lamprey Dicer has a longer IDR1 with a negative charge patch but this IDR1 is predicted to be outside of the RNA channel while lancelet Dicer IDR1 lacks the charge and is predicted to be outside (Fig. 1C). Apart from jawed vertebrates and sea urchin *Ciona intestinalis*, structure predictions of Dicer of deuterostomes and *Drosophila* have IDR1 located around the helicase domain but not inserted in the substrate channel (Fig. S1C).

**Figure 1.**
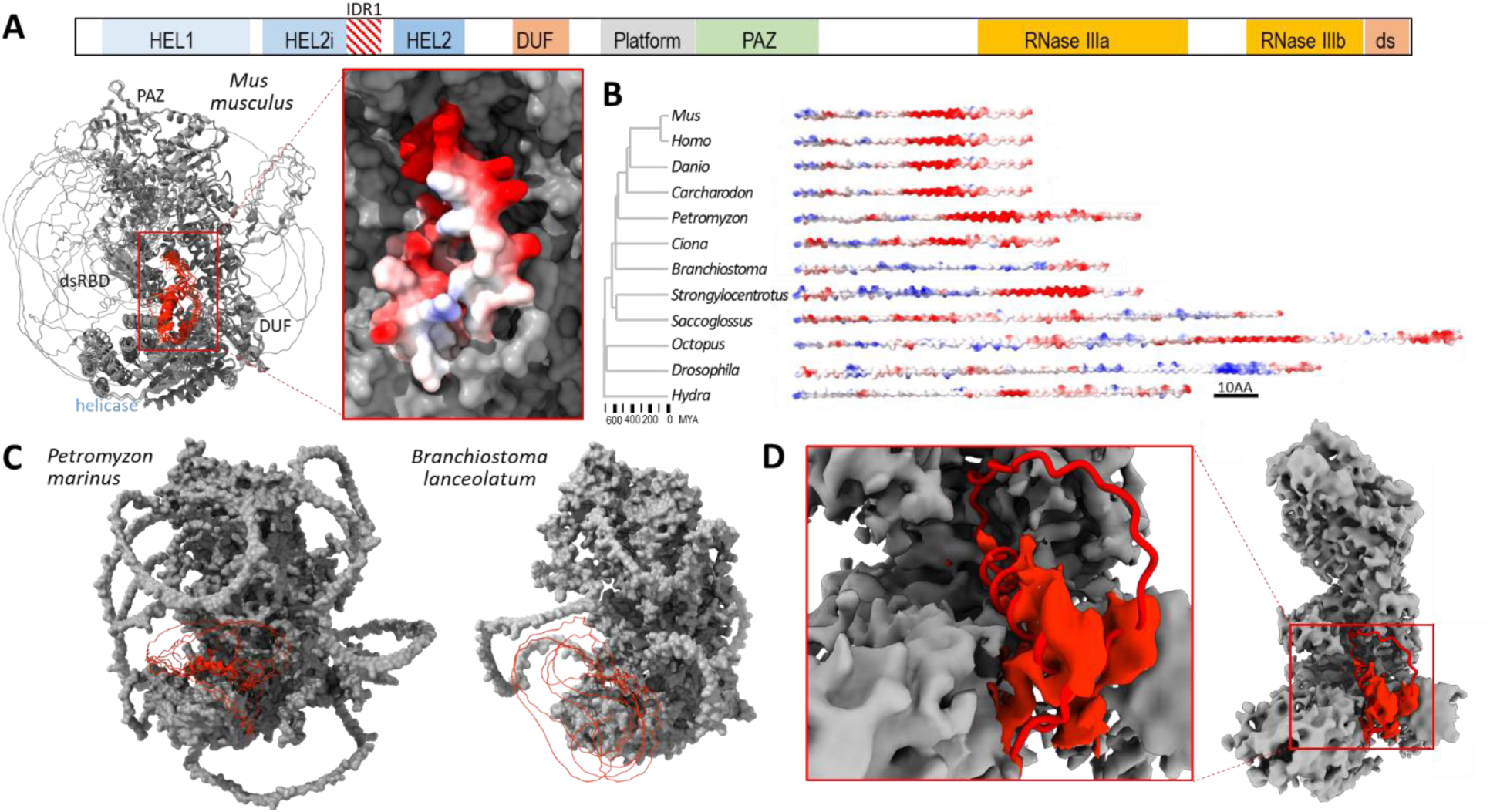
Vertebrate Dicers carry a negatively charged IDR in the helicase domain, which inserts into the RNA channel. (**A**) Schematic position of the IDR1 in Dicer and AlphaFold 3 (Abramson *et al*., 2024) prediction of the murine full-length Dicer structure where IDR1 is depicted in red color. Shown are the first five models aligned by the structured part of the enzyme. Image on the left shows one of the models where the negative charge patch of IDR1 (shown in red) faces the positively charged RNA channel. (**B**) Evolution of charged IDR1 in vertebrate Dicers. A schematic phylogenetic tree was obtained from TimeTree (Kumar *et al*, 2022), a schematic charge distribution on IDR1. From sharks (*Carcharodon*) to mammals is the IDR1 highly conserved. (**C**) AlphaFold 3 prediction of IDR1 positions in early branching chordates sea squirt *Ciona intestinalis* (IDR1 carries a negative charge patch) and lancelet *Branchiostoma lanceolatum* (IDR1 does not carry a charge). Five AlphaFold 3 predictions of IDR1 as red ribbon models were superimposed on one AlphaFold 3 model of the rest of Dicer, shown as a gray surface model. (**D**) A ribbon model of IDR1 overlap with cryo-EM data (in grey) (Lee *et al*., 2023) suggests an extra unassigned density (in red) in cryo-EM data matches the base of IDR1. UCSF ChimeraX (Meng *et al*., 2023) was used for all structural visualizations.

We also reviewed published cryo-EM data (Lee *et al*., 2023; Zapletal *et al*., 2022) to assess if any unassigned density could be observed in the area predicted to accommodate IDR1 in Dicer. This analysis revealed density overlapping with the AlphaFold 3 model at the base of the IDR1 loop, partially matching an α-helix predicted in the IDR1 (Fig 1D). There was no clear density in the substrate channel, likely reflecting conformational flexibility of the remaining portion of the loop.

Altogether, predicted position, charge and IDR1 conservation pattern suggested that it may be another sequence element stabilizing the closed “L”-shaped conformation of Dicer similarly to the HEL1 subdomain (Zapletal *et al*., 2022). We thus tested the impact of the loss of IDR1 on Dicer conformations and small RNA biogenesis.

### Loss of IDR1 enhances dicing state formation

To determine how IDR1 contributes to conformational control of mammalian Dicer, we purified a recombinant mouse Dicer variant lacking IDR1 (Dicer^ΔIDR1^). The mutant protein eluted identically to the wild-type enzyme on a gel filtration column, indicating that removal of IDR1 does not perturb the overall folding or monomeric state of Dicer (Fig. S2). We then analyzed the structural states of Dicer^ΔIDR1^ in complex with pre-miR-15a substrate by electron cryo-microscopy (cryo-EM). We hypothesized that removal of this negatively charged region might destabilize the pre-dicing state and facilitate transition toward the dicing-competent conformation (Fig. 2A). Cryo-EM analysis of the Dicer^ΔIDR1^–pre-miR-15a complex resolved two distinct conformational states corresponding to the pre-dicing and dicing states (Fig. 2B, C). As established previously, the dicing state lacks density for the helicase domain (Fig. 2C), consistent with its disengagement from the mammalian Dicer core during the transition to catalysis (Lee *et al*., 2023; Zapletal *et al*., 2022). In contrast, wild-type mammalian Dicer bound to the Dicer–pre-miR-15a has been observed exclusively in the pre-dicing state under identical conditions (Zapletal *et al*., 2022). Importantly, particles assigned to the pre-dicing state were twice more populated that the particles adopting the dicing state and displayed a properly folded and well-resolved helicase domain, indicating that loss of IDR1 does not compromise helicase domain integrity. These observations indicate that IDR1 is not required for formation of the pre-dicing state *per se* but rather stabilizes this conformation and suppresses spontaneous progression toward catalysis. This is important for the formation of the triangular RNA–protein architecture, which represents a substrate-licensing checkpoint specific to authentic pre-miRNA.

**Figure 2.**
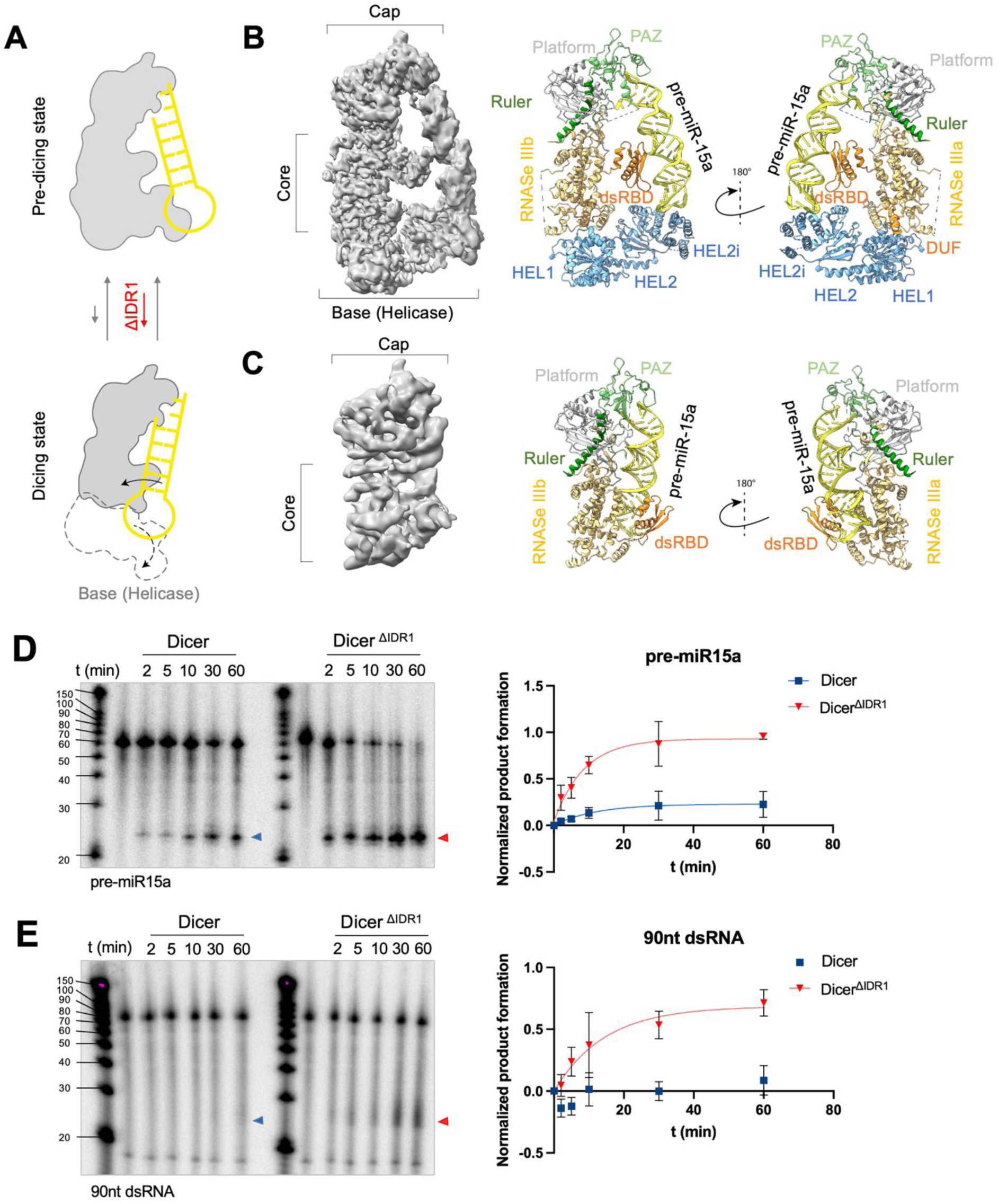
Structural analysis of Dicer^ΔIDR1^–pre-miR-15a complex (**A**) Structural models of the closed (pre-dicing) and open (dicing) conformations of Dicer, illustrating the proposed shift in conformational equilibrium upon IDR1 deletion. (**B**) Cryo-EM map (left) and corresponding ribbon representation (right) of the Dicer ^ΔIDR1^–pre-miR-15a complex in the pre-dicing state shown in two orientations rotated by 180°. (**C**) Same as in (B), showing the Dicer^ΔIDR1^–pre-miR-15a complex captured in the dicing state. (**D**) In vitro dicing assays and quantification comparing Dicer and Dicer ^ΔIDR1^ with pre-miR15a. (**E**) In vitro dicing assays and quantification comparing Dicer and Dicer ^ΔIDR1^ with 90nt dsRNA.

We also tested whether deletion of IDR1 alters Dicer catalytic activity *in vitro* using dicing assays with distinct RNA substrates. When assayed with pre-miR15a, Dicer^ΔIDR1^ exhibited a significantly increased cleavage efficiency compared with wild-type Dicer (Fig. 2D), in line with the enhanced sampling of the dicing state observed by cryo-EM. Wild-type Dicer showed very low activity toward a long 90-nt double-stranded RNA capped by a terminal loop (dsRNA), consistent with its suppression of RNA interference under physiological conditions (Flemr *et al*, 2013; Kennedy *et al*., 2015; Ma *et al*., 2008). Strikingly, deletion of IDR1 enhanced cleavage of a 90 nt dsRNA, indicating that the pre-dicing conformation stabilized by IDR1 normally acts as a barrier for canonical RNAi substrates (Fig. 2E). Together, these biochemical data corroborate the structural findings and demonstrate that IDR1 functions as a regulatory switch. This switch stabilizes the pre-dicing, substrate-licensing state of Dicer, thereby promoting high-fidelity miRNA processing while suppressing promiscuous cleavage of long double-stranded RNA.

### Loss of IDR1 activates RNAi

Increased abundance of dicing state of Dicer^ΔIDR1^ suggested that this Dicer variant might be efficiently supporting RNAi, similarly to the previously characterized ΔHEL1 variant naturally occurring in mouse oocytes (Demeter *et al*, 2019; Flemr *et al*., 2013; Zapletal *et al*., 2022). To test whether this truly occurs in mammalian cells, we used a canonical RNAi assay (Fig. S3A) in cultured cells transiently expressing Dicer variants. In mouse 3T3 *Pkr*^−/−^ cells, murine Dicer^ΔIDR1^ expression stimulated RNAi-mediated knock-down of a targeted luciferase reporter, albeit a bit less than Dicer^ΔHEL1^ (Fig. 3A). Remarkably, deletion of both, HEL1 and IDR1 showed an additive effect and resulted in even more efficient RNAi effect. The same result was obtained with human Dicer variants in HeLa *Pkr*^−/−^ cells suggesting the inhibitory effect on RNAi is conserved in mammals (Fig. 3B).

**Figure 3.**
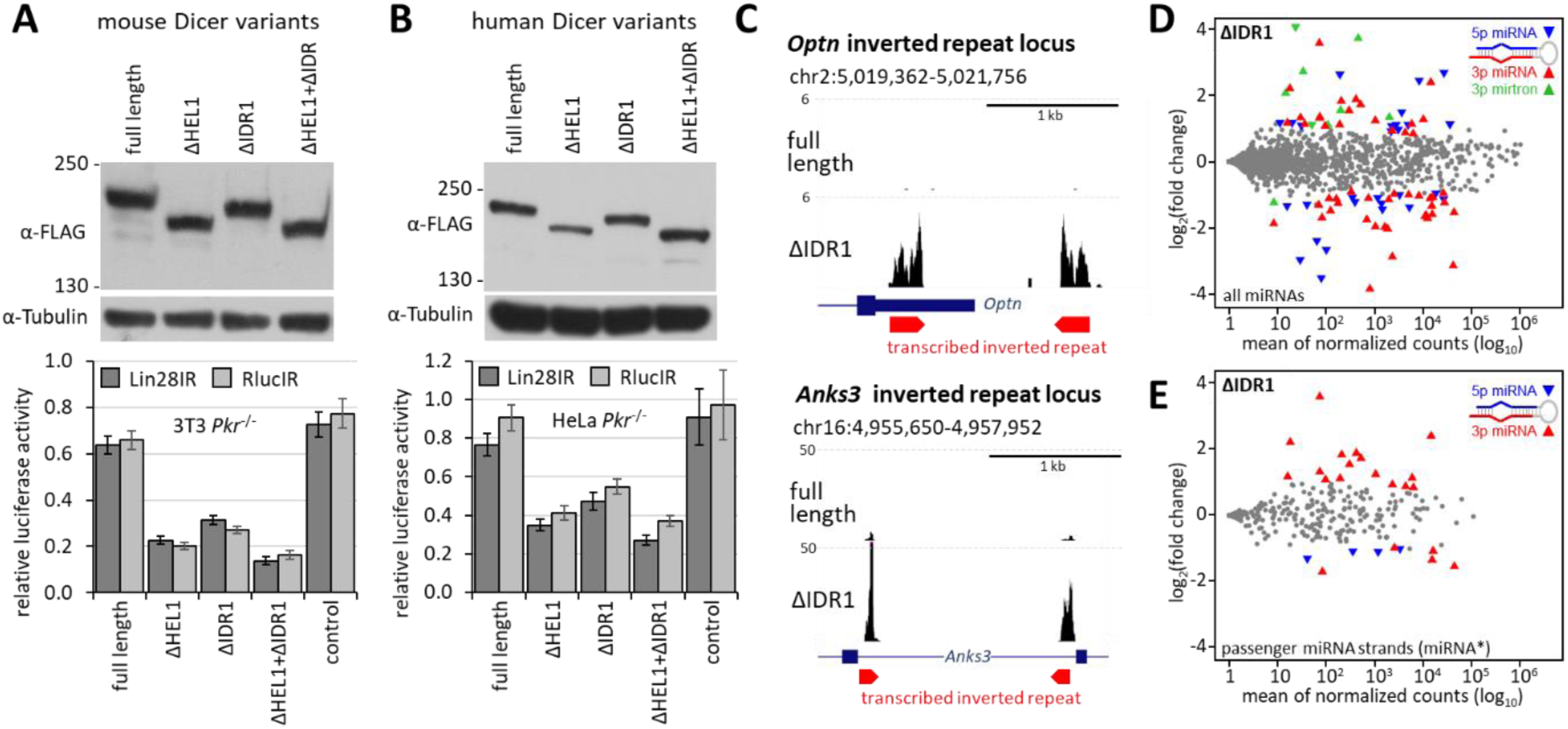
Functional analysis of ΔIDR1 Dicer in mammalian cells. (**A**) RNAi assay in murine 3T3 *Pkr*^−/−^ cells transiently expressing mouse C-terminally FLAG-tagged Dicer variants. Cells were co-transfected with a given Dicer variant, a plasmid expressing dsRNA (Lin28IR or RLucIR), a targeted *Renilla* luciferase reporter, and a non-targeted firefly luciferase reporter. As a control for Dicer effect was used a plasmid expressing LacZ (control). The scheme of the RNAi assay is shown in Fig. S3A. (**B**) RNAi assay in HeLa *Pkr*^−/−^ cells transiently expressing human Dicer variants. The organization of the assay is similar as in (A) except non-expressing plasmid was used as a control. Luciferase activity of the targeted reporter was normalized to a non-targeted firefly luciferase control and non-targeting dsRNA to account for sequence-independent effects and endogenous RNAi. (**C**) Stable ESC clones expressing only ΔIDR 1 Dicer variant exhibit strong increase in endogenous siRNA levels. Shown are UCSC browser snapshots of *Optn* and *Anks3* loci, which harbor transcribed inverted repeat sequences (red pentagons). Numbers at the y-scale represent counts per million of mapped AGO-bound small RNAs isolated from three different ESC clones for each Dicer variant. AGO-bound small RNAs were isolated using the TraPR approach (Grentzinger *et al*, 2020). (**D**) Ma plots depicting differentially expressed AGO-loaded miRNAs in ESCs expressing the full-length Dicer or its variant Dicer^ΔIDR1^. (**E**) Same analysis as in D but only passenger miRNA strand data are depicted.

To further examine how Dicer^ΔIDR1^ impacts small RNA biogenesis, we used *Dicer*^−/−^ mouse embryonic stem cell (ESC) (Murchison *et al*, 2005) to produce ESC clones stably expressing full-length Dicer or Dicer^ΔIDR1^ variants (Fig. S3B) and sequenced their small RNAs associated with Argonaute proteins. Inspection of small RNAs mapped to *Optn* and *Anks3*, two loci known to generate long dsRNA hairpins, showed strong upregulation of 21-23 small RNAs, consistent with increased efficiency of processing of long dsRNA into siRNAs by Dicer^ΔIDR1^ variant (Fig. 3C). These results are thus consistent with structural and *in vitro* experiments and reveal that IDR1 has an autoinhibitory role, which is suppressing RNAi in parallel to the previously discovered autoinhibitory role of HEL1. Disruption of either of these two elements is sufficient to activate RNAi while the maximum stability of the pre-dicing state is achieved when both elements are present.

### IDR1 is required for fidelity of miRNA biogenesis

Analysis of small RNAs in ESCs expressing Dicer^ΔIDR1^ also revealed dysregulation of multiple miRNAs, which showed features observed previously in ESCs expressing Dicer^ΔHEL1^ (Zapletal *et al*., 2022): increased levels of multiple 3p passenger strands and mirtrons (Fig. 3D). Mirtrons are non-canonical miRNAs whose stem loop precursors are formed by splicing of small introns (Berezikov *et al*, 2007) and have longer stems, which make them suboptimal substrates for normal Dicer while Dicer^ΔHEL1^ cleaves them more efficiently (Zapletal *et al*., 2022). We have found that mirtrons, showing increased abundance in ESCs expressing Dicer^ΔHEL1^ (Zapletal *et al*., 2022) also had higher levels in Dicer^ΔIDR1^ expressing ESCs (Fig. S3).

To further delineate the role of IDR1, we have produced ESC clones expressing additional Dicer variants where IDR1 was mutated to reduce the negative charge or replaced with IDRs of similar lengths from HSP90 and PAPD7 proteins (Fig. 4A, B and S4A). Sequencing of small RNAs confirmed increased production of siRNAs (Fig. S4B) and showed dysregulation of miRNAs in all Dicer variants (Fig. 4C, D). While all variants showed increased levels of mirtrons (Fig. 4D) and a subset of 3p passenger strands (Fig. S4B), the mutated Dicer variant where the distal charge of IDR1 was mutated to alanines showed a mild effect. Apparently, just neutralizing a large part of the charge was not sufficient to disrupt the role of IDRI completely.

**Figure 4.**
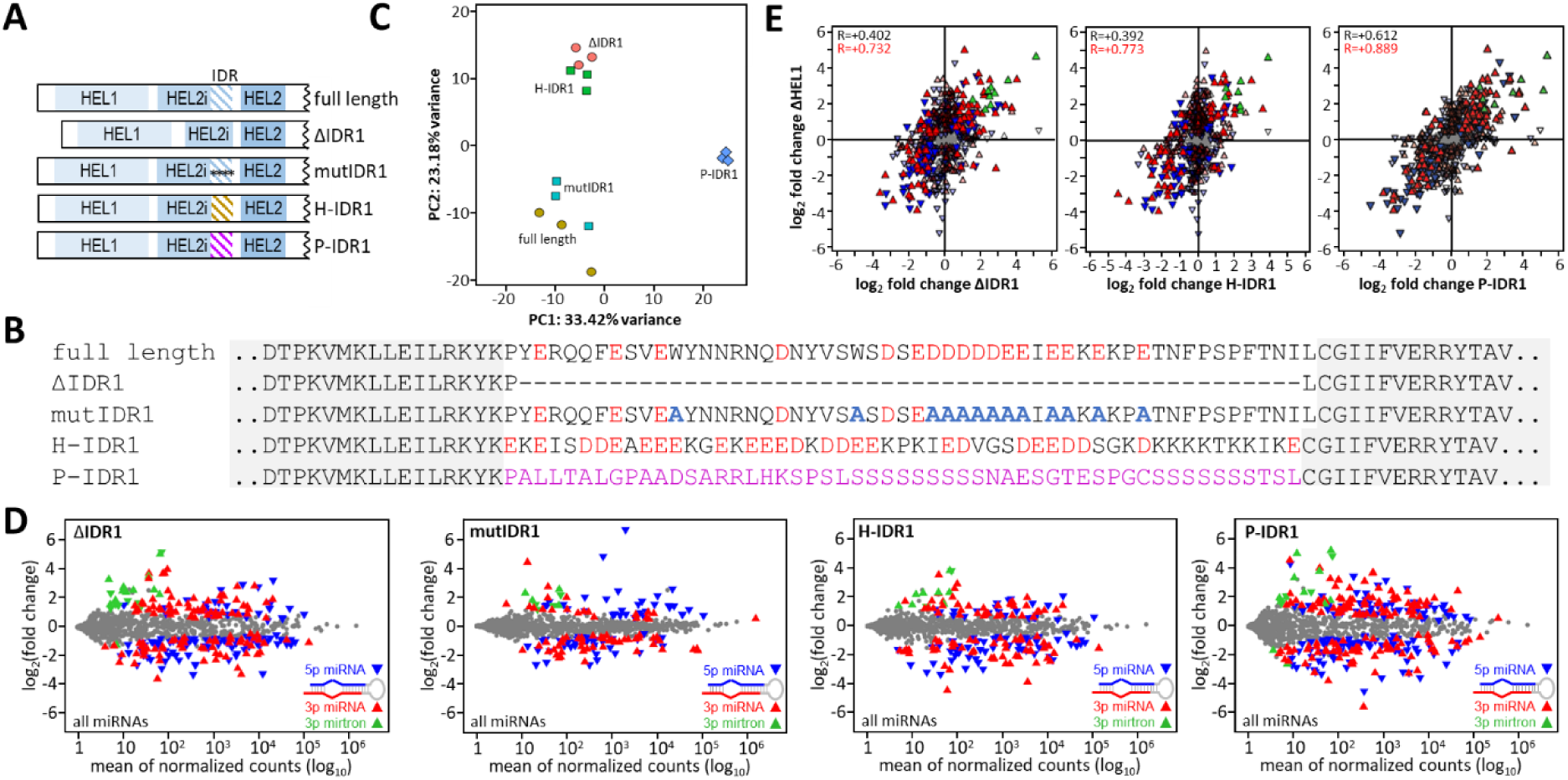
miRNA biogenesis in IDR1 mutants is affected similarly to the loss of the HEL1 domain. (A) Schematic depiction of Dicer variants used for experiments. (B) Sequences of IDR1 in Dicer variants shown in (A). H-IDR1 = HSP90ab IDR sequence, P-IDR1 = PAPD7 IDR sequence. (C) PCA analysis of RNA-seq of ESC clones (three ESC clones with each Dicer variant). (D) MA plots depicting differential miRNA expression in mutant ESC clones relative to clones expressing the full-length Dicer. (E) Correlation of relative miRNA changes in Dicer IDR mutants and changes in ESCs expressing Dicer^ΔHEL1^ (Zapletal *et al*., 2022).

Altogether, these data suggest that IDR1 fulfils a role analogous to HEL1 – IDR1 stabilizes the pre-dicing state, thus faithful recognition of pre-miRNAs. Observed small RNA misregulation then combines different impacts of Dicer^ΔIDR1^ on small RNA biogenesis. Increased production of siRNAs and mirtrons are most likely linked to destabilized pre-dicing state resulting in more efficient accommodation of other substrates than normal-sized pre-miRNAs. At the same time, increased abundance of 3p passenger strands may reflect a shift in strand selection and Argonaute loading processes. The same pattern has been observed in mutants Dicer^ΔHEL1^ (Fig. S4D) where no defect in cleavage fidelity was observed *in vitro* (Zapletal *et al*., 2022). It is remarkable that miRNA changes in strong IDR mutants match miRNA changes previously observed in cells expressing Dicer^ΔHEL1^ (Fig. 4E and S4C,D). The main common factor is disengagement of the helicase domain from the mammalian Dicer core where the unlocked helicase becomes highly flexible – this increases dicing of suboptimal substrates such as mirtrons and long dsRNA and likely affects strand selection of a subset of miRNAs. We speculate that reengagement of the helicase domain with the core may be linked to the strand selection and Argonaute loading processes. In any case, IDR1 represents another mechanistical principle stabilizing pre-dicing state, which became conserved across jawed vertebrates. Ancestral protein reconstruction suggests the helicase domain lost its affinity to long dsRNA and ATPase activity earlier (Aderounmu *et al*, 2023), which implies that IDR1 is a more recent adaptation of vertebrate Dicers for high fidelity and selectivity of miRNA biogenesis.

## MATERIAL AND METHODS

### Cell culture and transfection

Mouse ESCs were cultured in 2i-LIF media: KnockOut-DMEM (ThermoFisher) supplemented with 15% fetal calf serum (Sigma), 1x L-Glutamine (Sigma), 1x non-essential amino acids (ThermoFisher), 50 µM β-Mercaptoethanol (ThermoFisher), 1000 U/mL LIF (Isokine), 1 µM PD0325901, 3 µM CHIR99021 (Selleck Chemicals), penicillin (100 U/mL), and streptomycin (100 µg/mL). All plastic was coated with 1% gelatin (Sigma) in PBS.

NIH 3T3 fibroblasts and HeLa cells were cultured in DMEM (Sigma) supplemented with 10% fetal calf serum, penicillin (100 U/mL), and streptomycin (100 µg/mL).

### RNAi activity in cultured cells assay

Effects of different Dicer isoforms on RNAi-mediated repression in *Pkr*^−/−^HeLa or 3T3 cells were monitored as described previously (Demeter *et al*., 2019). Briefly, cells were co-transfected with a plasmid expressing a Dicer variant (or LacZ as a negative control), dsRNA (Lin28IR, RlucIR, or MosIR), a targeted *Renilla* luciferase reporter with complementary sequences to dsRNA from Lin28IR and RlucIR, and a non-targeted firefly luciferase.

For transfection, cells were plated on 24-well plates, grown to 80% density and transfected using Lipofectamine 3000 (ThermoFisher) according to the manufacturer’s protocol. The total amount of transfected DNA was kept constant (1 µg/well).

Specific repression of the targeted *Renilla* luciferase was estimated as *Renilla* luciferase activity normalized to the non-targeted firefly luciferase activity, and non-specific effect of MosIR (expressing a non-targeting dsRNA). The value 1.0 corresponds to absence of RNAi, the value of LacZ negative control reflects repression mediated by endogenously-expressed Dicer.

### Western blotting

HeLa cells or ES cells transfected with Dicer variants were homogenized mechanically in RIPA lysis buffer supplemented with 2x protease inhibitor cocktail set (Millipore) and loaded with SDS dye. Protein concentration was measured by Bradford assay (Bio-Rad) and 80 μg of total protein was used per lane. Proteins were separated on 5.5% polyacrylamide (PAA) gel and transferred on PVDF membrane (Millipore) using semi-dry blotting for 50 min, 35 V. The membrane was blocked in 5% skim milk in TBS-T, Dicer was detected using anti-FLAG (M2 mouse monoclonal antibody, Sigma #F3165, dilution 1:2,000) and incubated overnight at 4°C. Secondary HRP-conjugated anti-Mouse IgG binding protein (Santa-Cruz #sc-525409, dilution 1:50,000) or anti-Mouse-HRP antibody (Bio-Rad #1706516, dilution 1:100,000) was incubated 1 h at room temperature. For TUBA4A detection, samples were run on 10% PAA gel and incubated overnight at 4 °C with anti-Tubulin (Sigma, #T6074, dilution 1:10,000) mouse primary antibody. HRP-conjugated anti-mouse IgG binding protein (Santa-Cruz, #sc-525409, dilution 1:50,000) was used for detection. Signal was developed on films (X-ray film Blue, Cole-Parmer #21700-03) using SuperSignal West Femto Chemiluminescent Substrate (Thermo Scientific).

### Small RNA sequencing

ESCs were washed with PBS, homogenized in Qiazol lysis reagent (Qiagen), and total RNA was isolated by the phenol-chloroform extraction and ethanol precipitation method (Toni *et al*, 2018). After isolation, RNA quality was verified by electrophoresis in 1% agarose gel. For analysis of AGO-associated small RNAs, ESCs were used for isolation of RISC complexes by Trans-kingdom, rapid, affordable Purification of RISCs (TraPR - Lexogen) according to manufacturer’s instructions. RISC-associated small RNAs were extracted by a mixture of acidic phenol, chloroform, and isoamyl alcohol and precipitated by cold isopropanol. RISC-associated small RNAs were immediately used for small RNA library preparation.

Small RNA libraries were prepared using the NEXTFLEX® Small RNA-Seq Kit v4 for Illumina and Element® platforms (Revvity) according to the manufacturer’s protocol. 3′ adapter ligation was performed at 25 °C for 1 hour, 13-16 cycles were used for PCR amplification and gel purification was performed for size selection. Libraries were separated in 2.5% agarose gel using 1× lithium borate buffer and visualized with ethidium bromide. The 150–160 bp product was cut off the gel and DNA was isolated using the MinElute Gel Extraction Kit (Qiagen). Final libraries were analyzed by an Agilent 2100 Bioanalyzer and sequenced by 75-nucleotide single-end reading using the Illumina NextSeq1000/2000 platform.

### Bioinformatic analyses

RNA-seq data were deposited to NCBI as BioProjects with accession numbers PRJNA1414948 and PRJNA1415121, a sample overview is provided in the Table S1

#### Mapping of small RNA-seq data

Small RNA-seq reads were trimmed using cutadapt version 2.5 (Martin, 2011). Next, NEXTflex adapters were trimmed. Additionally, the N-nucleotides on ends of reads were trimmed and reads containing more than 10% of the N-nucleotides were discarded:

cutadapt --format=“fastq” --front=”TCTTTCCCTACACGACGCTCTTCCGATCT “ --adapter=”TGGAATTCTCGGGTGCCAAGG “ --error-rate=0.075 --times=2 --overlap=14 --minimum-length=12 --max-n=0.1 --output=”${TRIMMED}.fastq” --trim-n --match-read-wildcards ${TMP}.fastq

Trimmed reads were mapped to the mouse (mm10) genome using STAR aligner (Dobin *et al*, 2013) with following parameters:

STAR --readFilesIn ${TRIMMED}.fastq.gz --runThreadN 4 –genomeDir ${GENOME_INDEX} --genomeLoad LoadAndRemove --readFilesCommand unpigz -c --readStrand Unstranded --limitBAMsortRAM 20000000000 --outFileNamePrefix ${FILENAME} --outReadsUnmapped Fastx --outSAMtype BAM SortedByCoordinate --outFilterMultimapNmax 99999 --outFilterMismatchNoverLmax 0.1 --outFilterMatchNminOverLread 0.66 -- alignSJoverhangMin 999 --alignSJDBoverhangMin 999

#### miRNA expression analyses

Mapped reads were counted using program featureCounts (Liao *et al*, 2014). Only reads with lengths 19-25nt were selected from the small RNA-seq data:

`featureCounts -a ${ANNOTATION_FILE} -F ${FILE} -minOverlap 15 - fracOverlap 0.00 -s 1 -M -O -fraction -T 8 ${FILE}.bam`

The miRBase 22.1. (Kozomara *et al*, 2019) set of miRNAs was used for the annotation of small RNA-seq data for main figures, mirGeneDB annotation of high-confidence miRNAs^55^ was used to make sure that results were not biased by annotated low-confidence miRNAs from the miRBase. Statistical significance and fold changes in gene expression were computed in R using the DESeq2 package^75^. Genes were considered to be significantly up- or down-regulated if their corresponding p-adjusted values were smaller than 0.05.

#### miRNA expression plots – normalization of data, miRNA and miRNA* sorting

The annotation of the mirtrons was taken from Ladewig et al. (Ladewig *et al*, 2012). mirGeneDB annotation of high-confidence miRNAs (Clarke *et al*, 2025) was used to make sure that results were not biased by annotated low-confidence miRNAs from the miRBase. The DESeq2 baseMean and fold changes were plotted and visualized by home-made R scripts. The MA plots related to the dominant or passenger strand miRNAs contain only the corresponding miRNAs, all the miRNAs otherwise.

### Preparation of recombinant proteins for structural studies

Purification was carried out following protocols described in (Zapletal *et al*., 2022) with minor modifications. The coding sequence of the mouse Dicer^ΔIDR1^ variant, along with the required regulatory sequences was transposed into bacmid using *E. coli* strain DH10bac. Bacmid was transfected into the *Sf9* cells using FuGENE Transfection Reagent (Eastport) and the resulting viral particles were amplified in the culture. Dicer^ΔIDR1^ variant was expressed in 200 ml of Hi5 cells infected (at 1.2×10^6^ cells/ml) with the corresponding P1 virus at multiplicity of infection >1. The cells were harvested 48 hours post infection, washed by 1x PBS, and stored at -80°C. All further steps were carried out at + 4°C. Cell pellets were resuspended in 30 ml of cold lysis buffer containing 50 mM Tris (pH 8.0), 300 mM NaCl, 0.4% Triton X-100, 10% (v/v) glycerol, 10 mM imidazole, 1 mM DTT, 2 mM MgCl_2_, benzonase (250U), and protease inhibitors (0.66 μg/ml pepstatin, 5 μg/ml benzamidine, 4.75 μg/ml leupeptin, 2 μg/ml aprotinin) (Applichem). The cells were gently shaken for 10 min at 4°C and then briefly sonicated. The lysate was centrifugated at 21,000xg for 1 hour at 4°C. The supernatant was passed through a column containing 2.5 ml NiNTA-agarose (QIAGEN). The affinity matrix was washed 2-times with 15 ml of washing buffer (50 mM Tris (pH 8.0), 500 mM NaCl, 1 mM DTT, 1 mM CaCl_2_, 10 mM imidazole and protease inhibitors) and 1-time with 15 ml of salt buffer (50 mM Tris (pH 8.0), 1500 mM NaCl, 1 mM DTT, 1 mM CaCl_2_, 10 mM imidazole and protease inhibitors) and finally washed once more with 15 ml of washing buffer. The protein was eluted two times with 3 ml of elution buffer (50 mM Tris (pH 8.0), 500 mM NaCl, 1 mM DTT, 1 mM CaCl_2_, 300 mM imidazole and protease inhibitors). The fractions containing protein were pooled and supplemented with 4 mM EDTA to chelate residual Mg^2+^ ions. After chelation, Dicer protein was concentrated to 1 ml using 100 kDa cut-off Vivaspin Turbo15 (Sartorius). The Dicer^ΔIDR1^ was further purified on a size exclusion column (Superose 6 Increase 10/300 GL, GE Healthcare) equilibrated with a buffer containing 50 mM Tris (pH 8.0), 150 mM NaCl, 1 mM DTT, 2 mM CaCl_2_ and protease inhibitors. Fractions containing the protein were pooled, concentrated, snap-frozen in liquid nitrogen, and stored at -80°C until further use.

### Preparation of recombinant proteins for *In vitro* cleavage assay

Dicer variants (wild type Dicer and Dicer^ΔIDR1^) used for *In vitro* cleavage assays were purified as described in the previous chapter, with the exception that the chelation step was omitted and all buffers were uniformly supplemented with 2 mM MgCl₂ instead of CaCl₂.

### *In vitro* cleavage assay

#### Substrate preparation

The RNA oligonucleotides were diluted to a concentration of 250 nM in nuclease-free water, after which they were mixed with T4 Polynucleotide Kinase buffer. To promote proper folding, the RNA was heated to 95°C for 3 min and then immediately cooled on ice for 5 min. Radiolabeling was carried out by adding RNase inhibitors (NEB), T4 polynucleotide kinase (NEB), and [γ-32P]-ATP (HARTMANN ANALYTIC). The mixture was incubated at 37°C for 10 min. The 5′-radiolabelled RNA was purified on G-25 columns (GE Healthcare) and diluted to a final concentration of 50 nM. The radiolabeled RNA Decade Marker (ThermoFisher Scientific) was prepared according to the manufacturer’s protocol. The RNA and the marker were aliquoted and stored at -20°C.

#### Nuclease-activity assay

Time-course reactions were carried out in a total volume of 10 μl containing 5 nM radiolabeled RNA substrate and 50 nM Dicer variant in buffer containing 30 mM Tris (pH 7.5), 30 mM NaCl, 1 mM DTT, and 2 mM MgCl₂. The reactions were incubated at 37 °C for 2, 5, 10, 30, or 60 min and terminated by addition of 10 μl of 95% formamide, followed by heating at 95 °C for 5 min. After electrophoresis on a denaturing 20% polyacrylamide gel containing 8 M urea, the gels were exposed for 10-15 hours onto a phosphor imaging screen (Fujifilm). The signal was detected using The Amersham Typhoon scanner (Cytiva) and analyzed in Multi Gauge v3.2 software.

The obtained band intensities, as background-corrected relative values (Multi Gauge v3.2 software), were quantified and normalized in Prism. The signal from the no-enzyme control reaction was subtracted to correct for residual background. Normalized product formation was plotted as a function of time. The data are represented as the mean value for each time point, with standard deviation from four independent experiments. The data were fitted to a one-phase exponential association model using nonlinear regression.

### Cryo-EM specimen preparation and data acquisition

The complex was formed by direct mixing of Dicer^ΔIDR1^ and pre-miR15a at a 1:2 ratio in buffer containing 50 mM Tris (pH 8.0), 100 mM NaCl, 1 mM DTT, and 2 mM CaCl_2_. Final concentration of Dicer protein in solution was adjusted to about 2.5 uM and sample was incubated 30 min on ice. The QuantiFoil Cu 300 R1.2/1.3 grids were glow-discharged (110 seconds, hydrogen-oxygen) right before applying the cryo-EM specimen. In a Vitrobot Mark IV (ThermoFisher Scientific), 3.5 μl of the protein–RNA complex was applied on the grid from the plasma treated side. The grid was blotted for 6.0 sec, blot force -3, in 100% humidity at 4°C, and plunged in liquid ethane cooled by liquid nitrogen. The data were collected using Titan Krios (ThermoFirsher Scientific) transmission electron microscope operating at 300 kV with a Gatan K3 camera (Gatan). Images were recorded using SerialEM software (Mastronarde, 2005). The details about data acquisition, processing, structural refinement and validation are shown below.

Cryo-EM data collection and refinement statistics:

| Instrument |  |  |
| --- | --- | --- |
| Microscope | FEI TITAN KRIOS |  |
| Data collection |  |  |
| Sample | Dicer ΔIDR1–pre-miR-15a complex in pre-cleavage state | Dicer ΔIDR1–pre-miR-15a complex in dicing state |
| EMDB accession number | accession code pending | accession code pending |
| PDB accession number | accession code pending | accession code pending |
| Voltage (kV) | 300 |  |
| Detector (counting mode) | GATAN K3 (6k x 4k) |  |
| Symmetry | C1 |  |
| Electron dose (e<sup>-</sup>/ Å<sup>2</sup> ) | 58 |  |
| Defocus range (μm) | -1 to -2 |  |
| Pixel size (Å) | 0.51 |  |
| Movies collected | 15,764 |  |
| Initial particles | 2,933,476 |  |
| Final particles | 246,921 | 40,395 |
| **Model composition** |  |  |
| Protein residue | 1855 |  |
| **Refinement** |  |  |
| Unmasked/Corrected Map resolution (cryoSPARC)<br>0.143 (Å) | 4.0/3.2 | 6.7/4.3 |
| Map sharpening B-factor (Å<sup>2</sup>) | 92.5 | 156.9 |
| **R.m.s. deviation (PHENIX Refinement)** |  |  |
| Bond length (Å) | 0.002 | 0.002 |
| Bond angles (°) | 0.484 | 0.440 |
| B-factor (Å<sup>2</sup>) (protein) | 168.55 | 318.55 |
| B-factor (Å<sup>2</sup>) (nucleotide) | 276.29 | 403.02 |
| **Validation (MolProbity)** |  |  |
| MolProbity score | 1.35 | 1.31 |
| Clashscore | 4.34 | 3.74 |
| Poor rotamers (%) | 0.28 | 0.00 |
| Bad bonds (%) | 0.00 | 0.00 |
| Bad angles (%) | 0.00 | 0.00 |
| **Ramachandran plot** |  |  |
| Favored (%) | 97.25 | 97.18 |
| Allowed (%) | 2.75 | 2.82 |
| Disallowed (%) | 0.00 | 0.00 |

### Image processing of electron micrographs

All steps of cryo-EM data processing were performed using CryoSPARC (Punjani *et al*, 2017). The movies were first processed by Patch MotionCor2 (Zheng *et al*, 2017) for generation of motion corrected, dose-weighted micrograph stacks. The CTF parameters were estimated using Patch CTF Estimation and micrographs were further manually curated. For Dicer^ΔIDR1^–pre-miR15a complex a total of 2,933,476 particles were initially picked using the Blob Picker, of which 1,907,055 particles were extracted with a box size of 512 px and Fourier-cropped to 128 px. Following two rounds of 2D classification, particles corresponding to Dicer in multiple conformations were selected and subjected to heterogeneous refinement. Approximately 50% of these particles (262,495 particles) corresponded to the pre-dicing state, while ∼20% (101,158 particles) represented the dicing state. Subsequent processing included additional 2D classification, ab initio model generation, and 3D refinement steps (homogeneous refinement, non-uniform refinement, local CTF refinement, and local refinement). Particles were re-extracted with a box size of 512 px, Fourier-cropped to 256 px and processed using rounds of 2D classification, ab initio model generation, and 3D refinement steps. Using this workflow, 246,921 particles were obtained for the pre-dicing state, resulting in a reconstruction at 3.23 Å resolution, whereas 40,395 particles contributed to the dicing state reconstruction at a final resolution of 4.35 Å.

### Cryo-EM model building and refinement

The initial coordinates of the Dicer structures were obtained from the PDB database, specifically from the previously published work by (Zapletal *et al*., 2022). For the Dice^rΔIDR1^–pre-miR15a complex in pre-dicing state, structure 7YYM was used, and for the complex in the dicing state, structure 7ZPI was used. The coordinates were initially fitted into the corresponding density map using the ‘Fit in Map’ function in UCSF ChimeraX (Meng *et al*, 2023). The models’ fit within the maps was further improved using ISOLDE (Croll, 2018). The PDB coordinates and the density map were imported into the Coot and ‘The Real Space Refine Zone’ tool was used to optimize the fit of the coordinates within the map. Regions of low resolution and regions where the map lacked density were excluded from the structure. On the other hand, segments that were absent from the initial structures were built *de novo* into the density and then refined. The models were examined using Coot (Emsley *et al*, 2010) and ISOLDE with the following tools: ‘Density Analysis’, ‘Ramachandran Plot’, ‘Clashes’, ‘Rotamer Validation’, and ‘Peptide Bond Validation’. Further structural refinement was performed using Phenix and ISOLDE software (Liebschner *et al*, 2019). Finally, the overall refinement quality and structural statistics were assessed using MolProbity (Williams *et al*, 2018) and the validation tools provided by the Protein Data Bank (consortium, 2019).

### Data visualization

Molecular graphics were prepared using UCSF ChimeraX (Meng *et al*., 2023; Pettersen *et al*, 2021), developed by the Resource for Biocomputing, Visualization, and Informatics at the University of California, San Francisco.

## ACKNOWLEDGEMENTS

The main funding was provided by The Czech Ministry of Education, Youth and Sport (MEYS) OP JAK CZ.02.01.01/00/22_008/0004575. VB and DFJ were in part supported by PhD student fellowships from the Charles University; this work will be in part fulfilling requirements for a PhD degree as “school work”. Funding of DZ included OP RDE project „Internal Grant Agency of Masaryk University“ No.CZ.02.2.69/0.0/0.0/19_073/0016943. The Ministry of Education, Youth and Sports of the Czech Republic (MEYS CR) provided institutional support for CEITEC 2020 project LQ1601. We acknowledge Cryo-electron microscopy and tomography core facility of CIISB, Instruct-CZ Centre, supported by MEYS CR (LM2023042). Computational resources included e-Infrastruktura CZ (LM2023054, ID:90254) and ELIXIR-CZ (LM2018131, LM2023055, ID:90255) projects by MEYS CR.

## Author Contributions

Conceptualization: RS, PS

Investigation: DFJ, NN, RM, DZ, JPa, VB, DB, JPe, KK, PS, RS

Visualization: DFJ, NN, RM, DZ, JPa, RS, PS

Funding acquisition: RS, PS

Project administration: RM, RS, PS

Supervision: RM, KK, RS, PS

Writing – original draft: RS, PS

Writing – review & editing: DFJ, NN, RM, DZ, JPa, VB, DB, JPe, KK, PS, RS

## Declaration of interests

The authors declare no competing interests.

## SUPPLEMENTARY DATA

**Figure S1.**
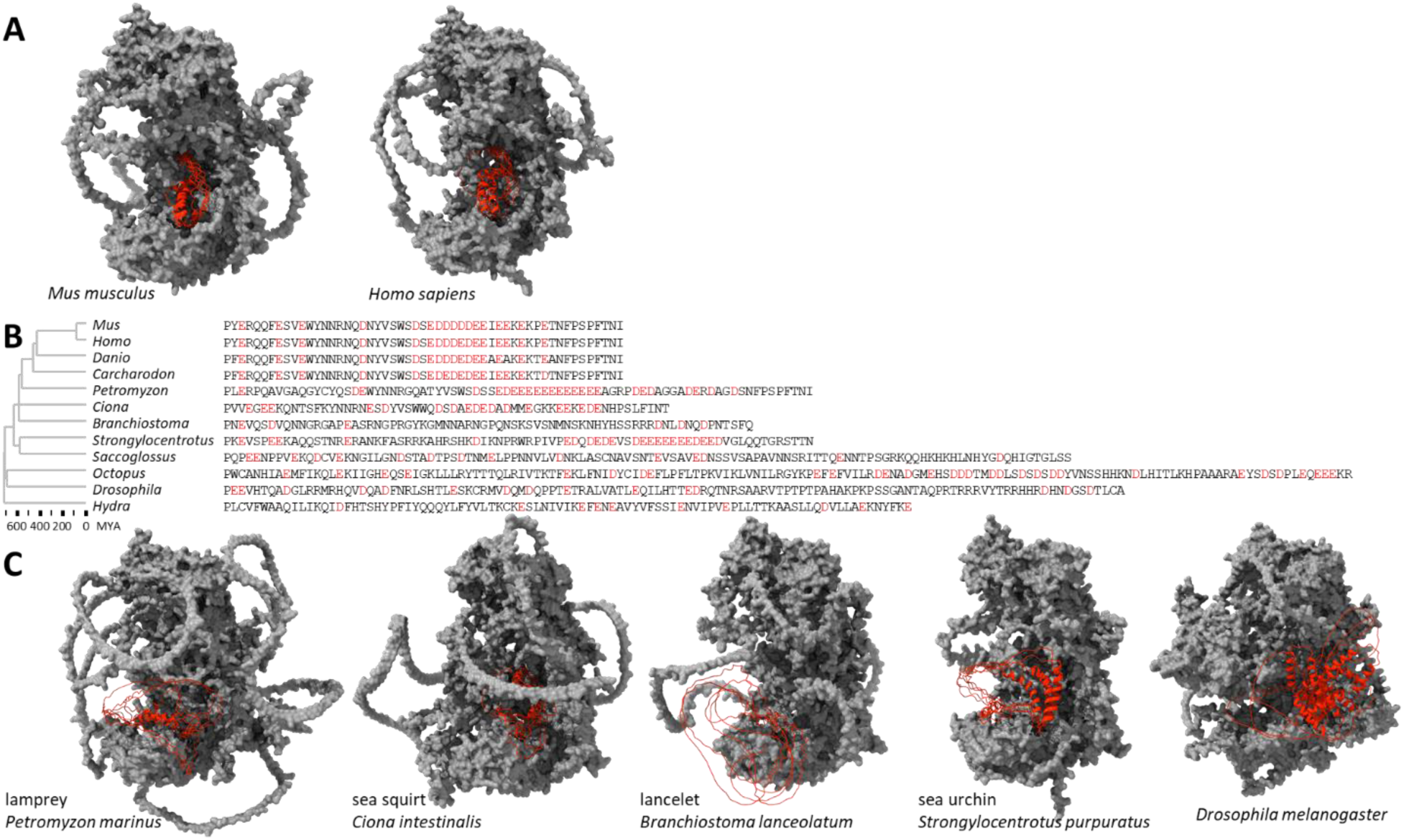
IDR1 in Dicer. (**A**) AlphaFold (Abramson *et al*., 2024) prediction of the murine and human full-length Dicer structures where IDR1 is depicted in red color. Five AlphaFold predictions of IDR1 as red ribbon models were superimposed on one AlphaFold model of the rest of Dicer, shown as a gray surface model. (**B**) IDR1 sequences of a set of Dicer proteins, negatively charged amino acids are shown in red font. (**C**) AlphaFold models of IDRs in selected phylogenetically more distant animals. Models were constructed as described in above.

**Figure S2.**
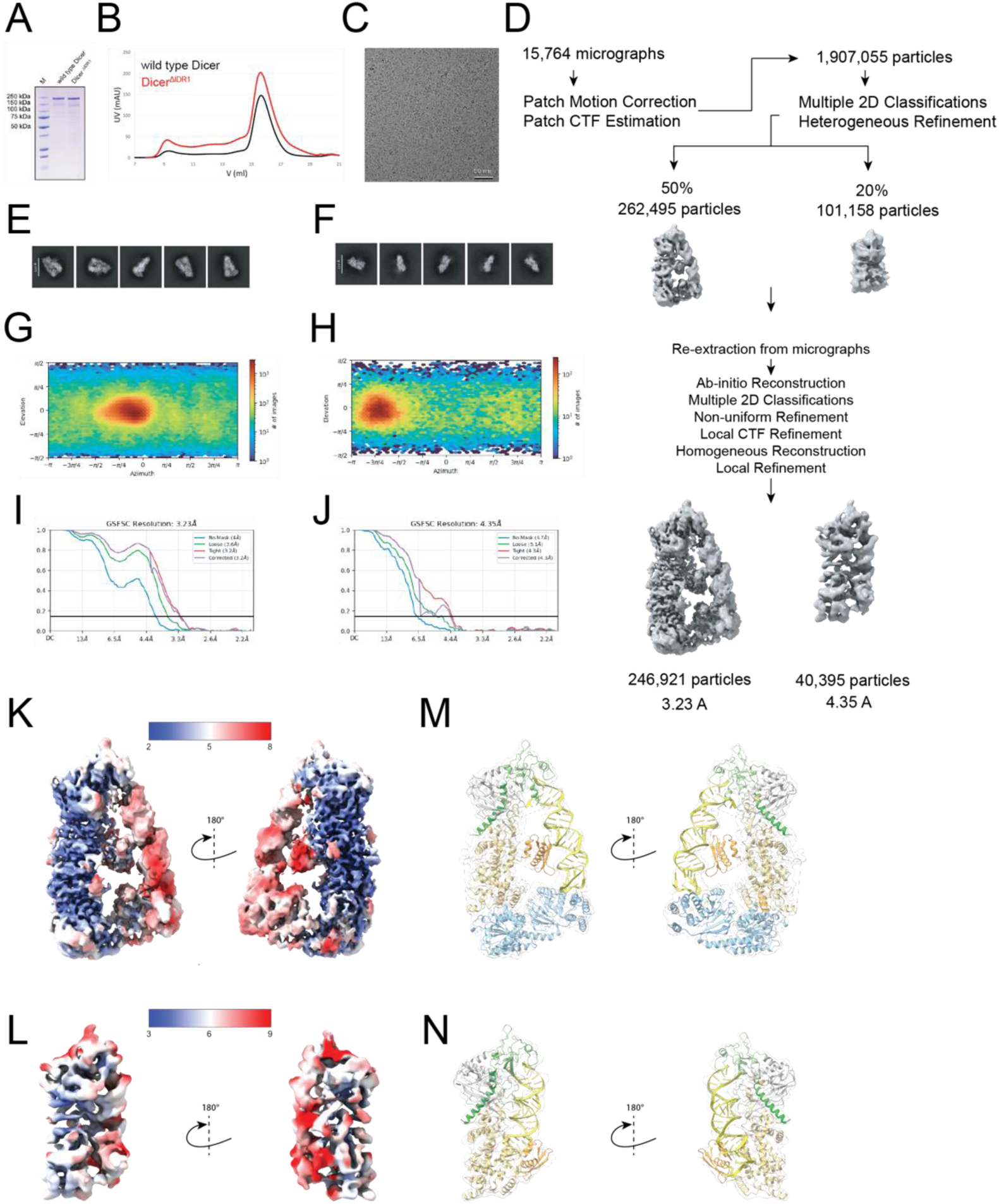
Reconstitution and cryo-EM analyses of Dicer ^ΔIDR1^–pre-miR-15a complex. (**A**) SDS-page analysis of wild type Dicer and Dicer ^ΔIDR1^. (**B**) Gel filtration analysis of wild type Dicer and Dicer ^ΔIDR1^. (**C**) Representative cryo-EM micrograph of the Dicer ^ΔIDR1^–pre-miR-15a complex. (**D**) Outline of the image processing steps used to obtain the 3.23-Å-resolution cryo-EM reconstruction of the Dicer–pre-miR-15a complex in pre-dicing state and 4.35-Å-resolution cryo-EM reconstruction of the Dicer–pre-miR-15a complex in dicing state. (**E-F**) Representative 2D class averages for the pre-dicing state (**E**) and dicing state (**F**). (**G-H**) Heat map for distribution of particles for the final 3D reconstruction of pre-dicing state (**G**) and dicing state (**H**). (**I-J**) FSC curves and resolutions calculated in cryoSPARC during final refinement for pre-dicing state (**I**) and dicing state (**J**). (**K-L**) Local resolution map of the final 3D reconstruction for pre-dicing state (**K**) and dicing state (**L**). (**M-N**) Structural model superposed to final cryo-EM density for pre-dicing state (**M**), dicing state (**N**).

**Figure S3.**
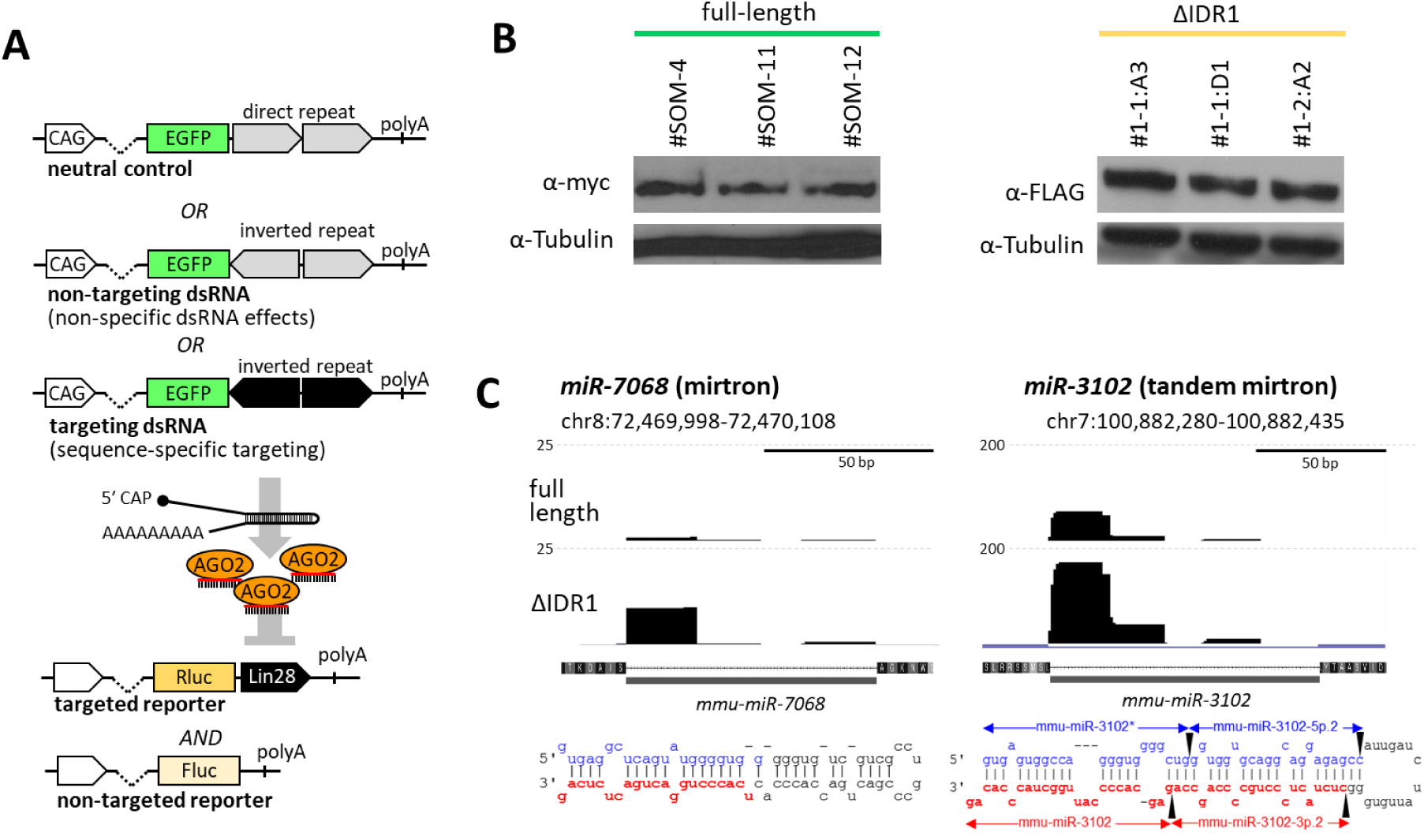
Dicer lacking IDR1 supports RNAi. (**A**) A schematic overview of the luciferase-based RNAi assay. (**B**) initial characterization of ESC clones expressing full-length Dicer and Dicer^ΔIDR1^ variant. (**C**) Increased abundance of mirtrons in ESCs expressing Dicer^ΔIDR1^. Shown are changes in expression of two highly upregulated mirtrons, which have extended stems relative to canonical miRNA precursors and were also upregulated in ESCs expressing Dicer^ΔIDR1^ (Zapletal *et al*., 2022). Above precursor schemes are shown UCSC browser snapshot tracks of small RNA-seq using the TraPR column (Grentzinger *et al*., 2020); the vertical scale is in CPMs. pre-miRNA folds were adopted from miRBase (Kozomara *et al*., 2019)). The miR-3102 mirtron is unique as its extended stem is cleaved by Dicer twice to release two miRNAs.

**Figure S4.**
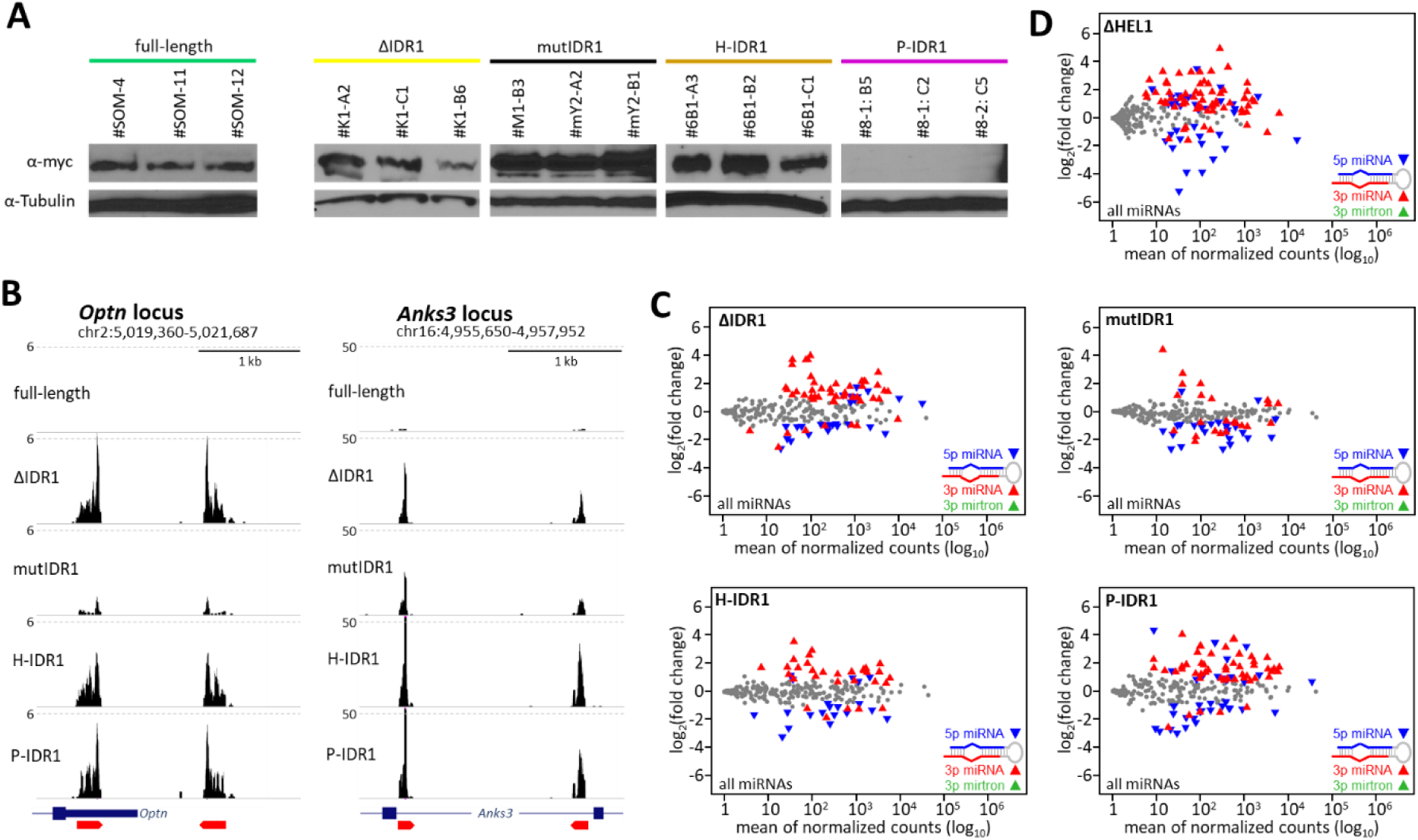
Small RNA changes in ESC lines expressing IDR1 mutants. (**A**) Dicer expression in different clones detected by Western blotting. P-IDR1 clones have very low Dicer expression but it is apparently sufficient for miRNA biogenesis. (**B**) Endogenous siRNA loci depicted in Fig 3B. Shown are UCSC browser snapshots of *Optn* and *Anks3* loci, which harbor transcribed inverted repeat sequences (red pentagons) and small RNAs 19-32 nucleotides mapping to these loci. (**C**) MA plots selectively depicting passenger strands reveal preferential upregulation of a subset of 3p strands (whose 5′ end is produced by Dicer cleavage), which is a pattern observed previously in (**D**) Dicer^ΔIDR1/ΔIDR1^ ESCs (Zapletal *et al*., 2022).

**Table S1.** Small RNA-seq sample overview.

| experiment | library | stage | sample type | genotype | clone |
| --- | --- | --- | --- | --- | --- |
| PRJNA1414948 | SAMN54917528 | mouse ESC | total small RNA | ΔIDR I | #K1-A2 |
| PRJNA1414948 | SAMN54917529 | mouse ESC | total small RNA | ΔIDR I | #K1-B6 |
| PRJNA1414948 | SAMN54917530 | mouse ESC | total small RNA | **ΔIDR I** | **#K1-C1** |
| PRJNA1414948 | SAMN54917531 | mouse ESC | total small RNA | full length | #SOM-4 |
| PRJNA1414948 | SAMN54917532 | mouse ESC | total small RNA | full length | #SOM-11 |
| PRJNA1414948 | SAMN54917533 | mouse ESC | total small RNA | full length | #SOM-12 |
| PRJNA1414948 | SAMN54917534 | mouse ESC | total small RNA | H-IDR | Y#6B1-A3 |
| PRJNA1414948 | SAMN54917535 | mouse ESC | total small RNA | H-IDR | #6B1-B2 |
| PRJNA1414948 | SAMN54917536 | mouse ESC | total small RNA | H-IDR | #6B1-C1 |
| PRJNA1414948 | SAMN54917537 | mouse ESC | total small RNA | mutIDR | #M1-B3 |
| PRJNA1414948 | SAMN54917538 | mouse ESC | total small RNA | mutIDR | #mY2-A2 |
| PRJNA1414948 | SAMN54917539 | mouse ESC | total small RNA | mutIDR | #mY2-B1 |
| PRJNA1414948 | SAMN54917540 | mouse ESC | total small RNA | P-IDR | #8-1:B5 |
| PRJNA1414948 | SAMN54917541 | mouse ESC | total small RNA | P-IDR | #8-1:C2 |
| PRJNA1414948 | SAMN54917542 | mouse ESC | total small RNA | P-IDR | #8-2:C5 |
| PRJNA1415121 | SAMN54919159 | mouse ESC | total small RNA | ΔIDR I | #1-A2 |
| PRJNA1415121 | SAMN54919160 | mouse ESC | total small RNA | ΔIDR I | #1-A3 |
| PRJNA1415121 | SAMN54919161 | mouse ESC | total small RNA | ΔIDR I | #1-D1 |
| PRJNA1415121 | SAMN54919162 | mouse ESC | total small RNA | full length | #SOM-4 |
| PRJNA1415121 | SAMN54919163 | mouse ESC | total small RNA | full length | #SOM-11 |
| PRJNA1415121 | SAMN54919164 | mouse ESC | total small RNA | full length | #SOM-12 |
| PRJNA1415121 | SAMN54919165 | mouse ESC | TraPR | ΔIDR I | #1-A2 |
| PRJNA1415121 | SAMN54919166 | mouse ESC | TraPR | ΔIDR I | #1-A3 |
| PRJNA1415121 | SAMN54919167 | mouse ESC | TraPR | ΔIDR I | #1-D1 |
| PRJNA1415121 | SAMN54919168 | mouse ESC | TraPR | full length | #SOM-4 |
| PRJNA1415121 | SAMN54919169 | mouse ESC | TraPR | full length | #SOM-11 |
| PRJNA1415121 | SAMN54919170 | mouse ESC | TraPR | full length | #SOM-12 |

## Notes

### Competing Interest Statement

The authors have declared no competing interest.

## REFERENCES

Abramson J, Adler J, Dunger J, Evans R, Green T, Pritzel A, Ronneberger O, Willmore L, Ballard AJ, Bambrick J et al (2024) Accurate structure prediction of biomolecular interactions with AlphaFold 3. Nature 630: 493–500

Aderounmu AM, Aruscavage PJ, Kolaczkowski B, Bass BL (2023) Ancestral protein reconstruction reveals evolutionary events governing variation in Dicer helicase function. Elife 12

Bartel DP (2018) Metazoan MicroRNAs. Cell 173: 20–51

Berezikov E, Chung WJ, Willis J, Cuppen E, Lai EC (2007) Mammalian mirtron genes. Mol Cell 28: 328–336

Cenik ES, Fukunaga R, Lu G, Dutcher R, Wang Y, Tanaka Hall TM, Zamore PD (2011) Phosphate and R2D2 restrict the substrate specificity of Dicer-2, an ATP-driven ribonuclease. Mol Cell 42: 172–184

Clarke AW, Hoye E, Hembrom AA, Paynter VM, Vinther J, Wyrozemski L, Biryukova I, Formaggioni A, Ovchinnikov V, Herlyn H et al (2025) MirGeneDB 3.0: improved taxonomic sampling, uniform nomenclature of novel conserved microRNA families and updated covariance models. Nucleic Acids Res 53: D116–D128

consortium w (2019) Protein Data Bank: the single global archive for 3D macromolecular structure data. Nucleic Acids Res 47: D520–D528

Cordin O, Banroques J, Tanner NK, Linder P (2006) The DEAD-box protein family of RNA helicases. Gene 367: 17–37

Croll TI (2018) ISOLDE: a physically realistic environment for model building into low-resolution electron-density maps. Acta Crystallogr D Struct Biol 74: 519–530

Demeter T, Vaskovicova M, Malik R, Horvat F, Pasulka J, Svobodova E, Flemr M, Svoboda P (2019) Main constraints for RNAi induced by expressed long dsRNA in mouse cells. Life Sci Alliance 2

Dobin A, Davis CA, Schlesinger F, Drenkow J, Zaleski C, Jha S, Batut P, Chaisson M, Gingeras TR (2013) STAR: ultrafast universal RNA-seq aligner. Bioinformatics 29: 15–21

Emsley P, Lohkamp B, Scott WG, Cowtan K (2010) Features and development of Coot. Acta Crystallogr D Biol Crystallogr 66: 486–501

Fairman-Williams ME, Guenther UP, Jankowsky E (2010) SF1 and SF2 helicases: family matters. Curr Opin Struct Biol 20: 313–324

Flemr M, Malik R, Franke V, Nejepinska J, Sedlacek R, Vlahovicek K, Svoboda P (2013) A retrotransposon-driven Dicer isoform directs endogenous small interfering RNA production in mouse oocytes. Cell 155: 807–816

Grentzinger T, Oberlin S, Schott G, Handler D, Svozil J, Barragan-Borrero V, Humbert A, Duharcourt S, Brennecke J, Voinnet O (2020) A universal method for the rapid isolation of all known classes of functional silencing small RNAs. Nucleic Acids Res 48: e79

Jia H, Kolaczkowski O, Rolland J, Kolaczkowski B (2017) Increased Affinity for RNA Targets Evolved Early in Animal and Plant Dicer Lineages through Different Structural Mechanisms. Mol Biol Evol 34: 3047–3063

Jouravleva K, Golovenko D, Demo G, Dutcher RC, Hall TMT, Zamore PD, Korostelev AA (2022) Structural basis of microRNA biogenesis by Dicer-1 and its partner protein Loqs-PB. Mol Cell 82: 4049–4063 e4046

Jumper J, Evans R, Pritzel A, Green T, Figurnov M, Ronneberger O, Tunyasuvunakool K, Bates R, Zidek A, Potapenko A et al (2021) Highly accurate protein structure prediction with AlphaFold. Nature 596: 583–589

Kennedy EM, Whisnant AW, Kornepati AV, Marshall JB, Bogerd HP, Cullen BR (2015) Production of functional small interfering RNAs by an amino-terminal deletion mutant of human Dicer. Proc Natl Acad Sci U S A

Ketting RF (2011) The many faces of RNAi. Dev Cell 20: 148–161

Ketting RF, Fischer SE, Bernstein E, Sijen T, Hannon GJ, Plasterk RH (2001) Dicer functions in RNA interference and in synthesis of small RNA involved in developmental timing in C. elegans. Genes Dev 15: 2654–2659

Kozomara A, Birgaoanu M, Griffiths-Jones S (2019) miRBase: from microRNA sequences to function. Nucleic Acids Res 47: D155–D162

Kumar S, Suleski M, Craig JM, Kasprowicz AE, Sanderford M, Li M, Stecher G, Hedges SB (2022) TimeTree 5: An Expanded Resource for Species Divergence Times. Mol Biol Evol

Ladewig E, Okamura K, Flynt AS, Westholm JO, Lai EC (2012) Discovery of hundreds of mirtrons in mouse and human small RNA data. Genome research 22: 1634–1645

Lau PW, Guiley KZ, De N, Potter CS, Carragher B, MacRae IJ (2012) The molecular architecture of human Dicer. Nat Struct Mol Biol 19: 436–440

Lau PW, Potter CS, Carragher B, MacRae IJ (2009) Structure of the human Dicer-TRBP complex by electron microscopy. Structure 17: 1326–1332

Lee YY, Lee H, Kim H, Kim VN, Roh SH (2023) Structure of the human DICER-pre-miRNA complex in a dicing state. Nature 615: 331–338

Liao Y, Smyth GK, Shi W (2014) featureCounts: an efficient general purpose program for assigning sequence reads to genomic features. Bioinformatics 30: 923–930

Liebschner D, Afonine PV, Baker ML, Bunkoczi G, Chen VB, Croll TI, Hintze B, Hung LW, Jain S, McCoy AJ et al (2019) Macromolecular structure determination using X-rays, neutrons and electrons: recent developments in Phenix. Acta Crystallogr D Struct Biol 75: 861–877

Liu Q, Rand TA, Kalidas S, Du F, Kim HE, Smith DP, Wang X (2003) R2D2, a bridge between the initiation and effector steps of the Drosophila RNAi pathway. Science 301: 1921–1925

Liu Z, Wang J, Cheng H, Ke X, Sun L, Zhang QC, Wang HW (2018) Cryo-EM Structure of Human Dicer and Its Complexes with a Pre-miRNA Substrate. Cell 173: 1191–1203 e1112

Ma E, MacRae IJ, Kirsch JF, Doudna JA (2008) Autoinhibition of human dicer by its internal helicase domain. J Mol Biol 380: 237–243

MacRae IJ, Zhou K, Li F, Repic A, Brooks AN, Cande WZ, Adams PD, Doudna JA (2006) Structural basis for double-stranded RNA processing by Dicer. Science 311: 195–198

Martin M (2011) Cutadapt removes adapter sequences from high-throughput sequencing reads. EMBnetJournal 17: 3

Mastronarde DN (2005) Automated electron microscope tomography using robust prediction of specimen movements. J Struct Biol 152: 36–51

Meng EC, Goddard TD, Pettersen EF, Couch GS, Pearson ZJ, Morris JH, Ferrin TE (2023) UCSF ChimeraX: Tools for structure building and analysis. Protein Sci 32: e4792

Murchison EP, Partridge JF, Tam OH, Cheloufi S, Hannon GJ (2005) Characterization of Dicer-deficient murine embryonic stem cells. Proc Natl Acad Sci U S A 102: 12135–12140

Pettersen EF, Goddard TD, Huang CC, Meng EC, Couch GS, Croll TI, Morris JH, Ferrin TE (2021) UCSF ChimeraX: Structure visualization for researchers, educators, and developers. Protein Sci 30: 70–82

Provost P, Dishart D, Doucet J, Frendewey D, Samuelsson B, Radmark O (2002) Ribonuclease activity and RNA binding of recombinant human Dicer. EMBO J 21: 5864–5874

Punjani A, Rubinstein JL, Fleet DJ, Brubaker MA (2017) cryoSPARC: algorithms for rapid unsupervised cryo-EM structure determination. Nat Methods 14: 290–296

Taylor DW, Ma E, Shigematsu H, Cianfrocco MA, Noland CL, Nagayama K, Nogales E, Doudna JA, Wang HW (2013) Substrate-specific structural rearrangements of human Dicer. Nat Struct Mol Biol 20: 662–670

Toni LS, Garcia AM, Jeffrey DA, Jiang X, Stauffer BL, Miyamoto SD, Sucharov CC (2018) Optimization of phenol-chloroform RNA extraction. MethodsX 5: 599–608

Welker NC, Maity TS, Ye X, Aruscavage PJ, Krauchuk AA, Liu Q, Bass BL (2011) Dicer’s helicase domain discriminates dsRNA termini to promote an altered reaction mode. Mol Cell 41: 589–599

Williams CJ, Headd JJ, Moriarty NW, Prisant MG, Videau LL, Deis LN, Verma V, Keedy DA, Hintze BJ, Chen VB et al (2018) MolProbity: More and better reference data for improved all-atom structure validation. Protein Sci 27: 293–315

Zapletal D, Kubicek K, Svoboda P, Stefl R (2023) Dicer structure and function: conserved and evolving features. EMBO Rep 24: e57215

Zapletal D, Taborska E, Pasulka J, Malik R, Kubicek K, Zanova M, Much C, Sebesta M, Buccheri V, Horvat F et al (2022) Structural and functional basis of mammalian microRNA biogenesis by Dicer. Mol Cell 82: 4064–4079 e4013

Zhang H, Kolb FA, Brondani V, Billy E, Filipowicz W (2002) Human Dicer preferentially cleaves dsRNAs at their termini without a requirement for ATP. EMBO J 21: 5875–5885

Zhang H, Kolb FA, Jaskiewicz L, Westhof E, Filipowicz W (2004) Single processing center models for human Dicer and bacterial RNase III. Cell 118: 57–68

Zheng SQ, Palovcak E, Armache JP, Verba KA, Cheng Y, Agard DA (2017) MotionCor2: anisotropic correction of beam-induced motion for improved cryo-electron microscopy. Nat Methods 14: 331–332

Zhu M, Yu X, Ahlberg PE, Choo B, Lu J, Qiao T, Qu Q, Zhao W, Jia L, Blom H et al (2013) A Silurian placoderm with osteichthyan-like marginal jaw bones. Nature 502: 188–193

